# Soil viral community dynamics over seven years of heat disturbance: spatial variation exceeds temporal in annually sampled soils

**DOI:** 10.1101/2024.05.27.596044

**Authors:** Samuel E. Barnett, Ashley Shade

## Abstract

Viruses are important components of the soil microbiome, influencing microbial population dynamics and the functions of their hosts. However, the relationships and feedbacks between virus dynamics, microbial host dynamics, and environmental disturbance is not understood. Centralia, PA, USA, is the site of an underground coal seam fire that has been burning for over 60 years. As the fire moves along the coal seam, previously heated soils cool to ambient temperature, creating a gradient of heat disturbance intensity and recovery. We examined annual soil viral population dynamics over seven consecutive years in Centralia using untargeted metagenome sequencing. Viral communities changed over time and were distinct between fire-affected and reference sites. Dissimilarity in viral communities was greater across sites (space) than within a site across years (time), and cumulative viral diversity more rapidly stabilized within a site across years than within a year across sites. There also were changes in CRISPR investment as soils cooled, corresponding to shifts in viral diversity. Finally, there were also differences in viral-encoded auxiliary metabolic genes between fire-affected and reference sites. These results indicate that despite high site-to-site soil viral diversity, there was surprising viral community consistency within a site over the years and shifting host-viral interactions in soils recovering from disturbance. Together, these results provide insights into how viral and host communities collectively respond to unpredicted environmental disturbance.

**Highlights:**

- In a seven-year annual study of temperate soils affected by an underground fire, viral communities displayed greater variability across sites (spatial) than over time (temporal).
- Viral communities were distinct between heated and unheated soils throughout sampling.
- Soil bacterial community composition correlated to viral community composition, though this relationship weakened when accounting for edaphic properties of soil temperature and pH.
- Soil viral communities may be less resilient to press disturbance than their host bacterial communities.
- Viral-host interactions may shift during soil recovery from long-term heating.

## 1 Introduction

Viruses are ubiquitous in the environment and form complex and dynamic relationships with their host populations. Viruses that infect bacteria or archaea, called phages, are of particular interest because they influence the dynamics and functions of their host populations (Weinbauer and Rassoulzadegan, 2004; Wigington et al., 2016; Liang et al., 2020; Roy et al., 2020; Jansson, 2023). Recent studies have begun to understand the roles phages play in complex natural habitats such as soils (Van Goethem et al., 2019; Trubl et al., 2020, 2021; Starr et al., 2021; Chevallereau et al., 2022; Muscatt et al., 2023). For example, phages have been shown to influence soil carbon and nitrogen cycling through mechanisms such as host population control, biomass turnover, and auxiliary metabolic genes (AMG) (Kuzyakov and Mason-Jones, 2018; Trubl et al., 2018, 2023; Starr et al., 2019; Braga et al., 2020; Barnett and Buckley, 2023; Tong et al., 2023). While it is clear that phages are essential components of ecosystems, we still have much to learn about how disturbances influence phage communities, especially disturbances related to human activity and climate change (Jansson and Wu, 2022; Liao et al., 2022; Santos-Medellín et al., 2023).

Disturbances can affect the population dynamics of soil phages and their host microorganisms (Voigt et al., 2021; Jansson and Wu, 2022). Pulse events, such as soil wetting, produce drastic shifts in the phage community structure, including increased phage abundance, particularly phages linked to hosts undergoing population growth (Van Goethem et al., 2019; Santos-Medellín et al., 2023). Press events, such as land use change, also significantly alter soil phage communities, and soil pH is an important determinant of phage composition across land use types (Liao et al., 2022). As phage population dynamics feedback on those of their hosts, their dynamics during and following disturbances could provide insights a microbiome’s overall recovery and stability. However, there are knowledge gaps in how soil phage communities shift during microbiome recovery from disturbances, particularly press disturbances (Jansson and Wu, 2022). The complex lifestyle of phages can make it difficult to predict their dynamics under changing environmental conditions. For example, phages can remain dormant for extended periods, as is expected in soils, as phages have been experimentally shown to persist for weeks in soils without a host (DiPietro et al., 2023).

We used the unique temperature disturbance gradient in Centralia, Pennsylvania, USA, to examine phage community diversity and 7-year dynamics as soils recover from a decades-old underground coal seam fire (Barnett and Shade, 2024a). The fires under Centralia have been burning along two roughly linear coal seams since 1962, elevating surface soil temperatures and releasing combustion products through soil vents (Elick, 2011). A disturbance gradient has formed as the fires advance along the coal seams. Yet undisturbed soils (reference) are in front of and alongside the fire front, disturbed heated soils (fire-affected) are directly above the active fire, and recovering soils are where the fire has already passed (Lee et al., 2017; Kearns and Shade, 2018; Barnett and Shade, 2024a). The disturbance gradient provides a window into the dynamics of microbial communities during an intense press disturbance and through their post-disturbance recovery. Our previous studies have demonstrated the bacterial community’s resilience to long-term heating in Centralia soils. The bacterial communities in the recovered soils were more similar to communities in reference soils than fire-affected soils (Lee et al., 2017). As fire-affected soils cooled over seven consecutive years, we further observed directional changes in the bacterial community structure, with the community becoming more similar to the unaffected reference communities over time (Barnett and Shade, 2024a).

No studies have examined soil viral dynamics under long-term soil heating conditions like those in Centralia. Furthermore, it is not clear how much soil viral communities change over multiple years. Given the high spatial variation in soil viral communities compared across soils that were meters to tens of meters apart (Santos-Medellín et al., 2022), it may be naively expected that phage communities should also be highly variable and inconsistent over time. We hypothesized that as Centralia soils cool and the residing bacterial communities recover (Barnett and Shade, 2024a), the phage populations will also stabilize. In other words, we expect the phage population dynamics to reflect parallel resilience to the bacterial populations they infect.

## 2 Materials and Methods

### 2.1 Study site and soil sampling

The Centralia study site has been described previously (Elick, 2011; Lee et al., 2017; Kearns and Shade, 2018; Barnett and Shade, 2024a). The sites in Columbia County, Pennsylvania, overlay an anthracite seam of the Buck Mountain coal bed that has been burning constantly since 1962. The fire lies about 46m below the surface, and heat and combustion gasses from the fire affect the surface, increasing surface soil temperatures and producing steam vents (Elick 2011). Based on our previous studies (Lee et al., 2017; Kearns and Shade, 2018; Barnett and Shade, 2024a), we selected ten sites for untargeted metagenome sequencing, including three reference sites and seven fire-affected sites (Fig. S1A, Table S1).

Soils were collected from each site from 2015 to 2021 in the first or second week of October. Twenty cm depth soil cores were collected as described (Barnett and Shade, 2024a), sieved at 4 mm, hand-homogenized, and flash-frozen in liquid nitrogen. *In situ,* soil, and air temperatures were collected with an electronic thermometer. Soil moisture was measured gravimetrically, and soil chemistry analysis was performed by the Soil and Plant Nutrition Lab at Michigan State University (East Lansing, USA; https://www.canr.msu.edu/spnl). As previously reported (Barnett and Shade, 2024a), soil temperature across the fire-affected sites decreased over the seven sampling years as the fire gradually advanced and those soils began to cool (Fig S1B).

### 2.2 DNA extraction and sequencing

DNA was extracted from 0.25 g of soil as previously described following the Griffiths et al. phenol-chloroform protocol (Griffiths et al., 2000), with the alternate use of 0.1 mm zirconium bead containing BeadBug homogenizer tubes (#Z763764; Benchmark Scientific, Sayreville USA) for bead beating. DNA was stored at −80°C until sequencing. Libraries were prepared and sequenced by the Research Technology Support Facility at Michigan State University (East Lansing, MI, USA) as previously described (Barnett and Shade, 2024b). At this stage, one sample (site Cen08 from 2019) was removed due to failed library preparation, and the remaining 69 samples were pooled. Metagenome sequencing was performed in two lanes of a NovaSeq S4 flow cell in 2×150bp paired-end format on a NovaSeq 6000 (Illumina, San Diego, USA). Raw sequencing data is available on the NCBI SRA (BioProject PRJNA974462).

### 2.3 Sequence processing

Metagenomic sequencing reads were processed as described (Barnett and Shade, 2024b) using a custom workflow based on the Joint Genome Institute (Clum et al., 2021). Quality-controlled reads were assembled within individual samples using SPAdes version v3.15.5 in metaSPAdes mode with assembler only (Nurk et al., 2017). Assemblies can be accessed through NCBI (BioProject PRJNA974462), and metagenome processing code is available via GitHub (https://github.com/ShadeLab/Centralia_7year_metagenome_processing_Barnett_2024).

All viral sequence processing code is available via GitHub (https://github.com/ShadeLab/Centralia_phages_Barnett). We used a custom workflow based on the VirSorter2 SOP (dx.doi.org/10.17504/protocols.io.bwm5pc86) to identify unique viral operational taxonomic units (vOTU; Fig. 1). We first ran virsorter2 version 2.2.4 (Guo et al., 2021), removing contigs less than 5000 bp, keeping the original sequences, and using a minimum score of 0.5. The identified contigs were assessed with CheckV version 1.0.1 (Nayfach et al., 2021) to identify viral and host genes and trim sequences to viral regions. These CheckV-trimmed sequences, including both viral and proviral sequences, were then again run through virsorter2 but this time without filtering (--viral-gene-enrich-off and --provirus-off) to generate files utilized by DRAM-v. Finally, these sequences were annotated with DRAM-v. The sequences were then categorized into two “keep” categories (cat 1: number of viral genes > 0, and cat 2: number of viral genes = 0 and number of host genes = 0 or score >= 0.95 or number of hallmark genes >2), or a manual curation category (number of viral genes = 0 and number of host genes = 1 and length >= 10 Kbp). All other sequences were removed. Sequences in the keep category 2 that contained genes common in both viruses and hosts and can lead to false positive assignment were moved into the manual curation category. These suspect annotations were “carbohydrate kinase,” “carbohydrate-kinase,” “glycosyltransferase,” “glycosyl transferase,” “glycosyl transferaseendonuclease,” “nucleotide sugar epimerase,” “nucleotide sugar-epimerase,” “nucleotide-sugar epimerase,” “nucleotide-sugar-epimerase,” “nucleotidyltransferase,” “nucleotidyl transferase,” “nucleotidyl-transferase,” “plasmid stability,” “endonuclease”. The manual curation category was then further filtered to those in one of two further categories. The first category included sequences having structural genes, hallmark genes, depletion in annotations, or enrichment for hypotheticals. The second category included sequences lacking hallmarks, but over 50% of the annotated genes hit a virus, and at least half of those have viral bitscore >100, and the contig is less than 50kb in length. Final viral sequences from all samples that passed filtering were then binned into vOTU using the MMseqs2 version 14.7e284 (Steinegger and Söding, 2017) easy-cluster tool with a minimum sequence identity of 0.95 and target coverage of 0.85.

**Figure 1:**
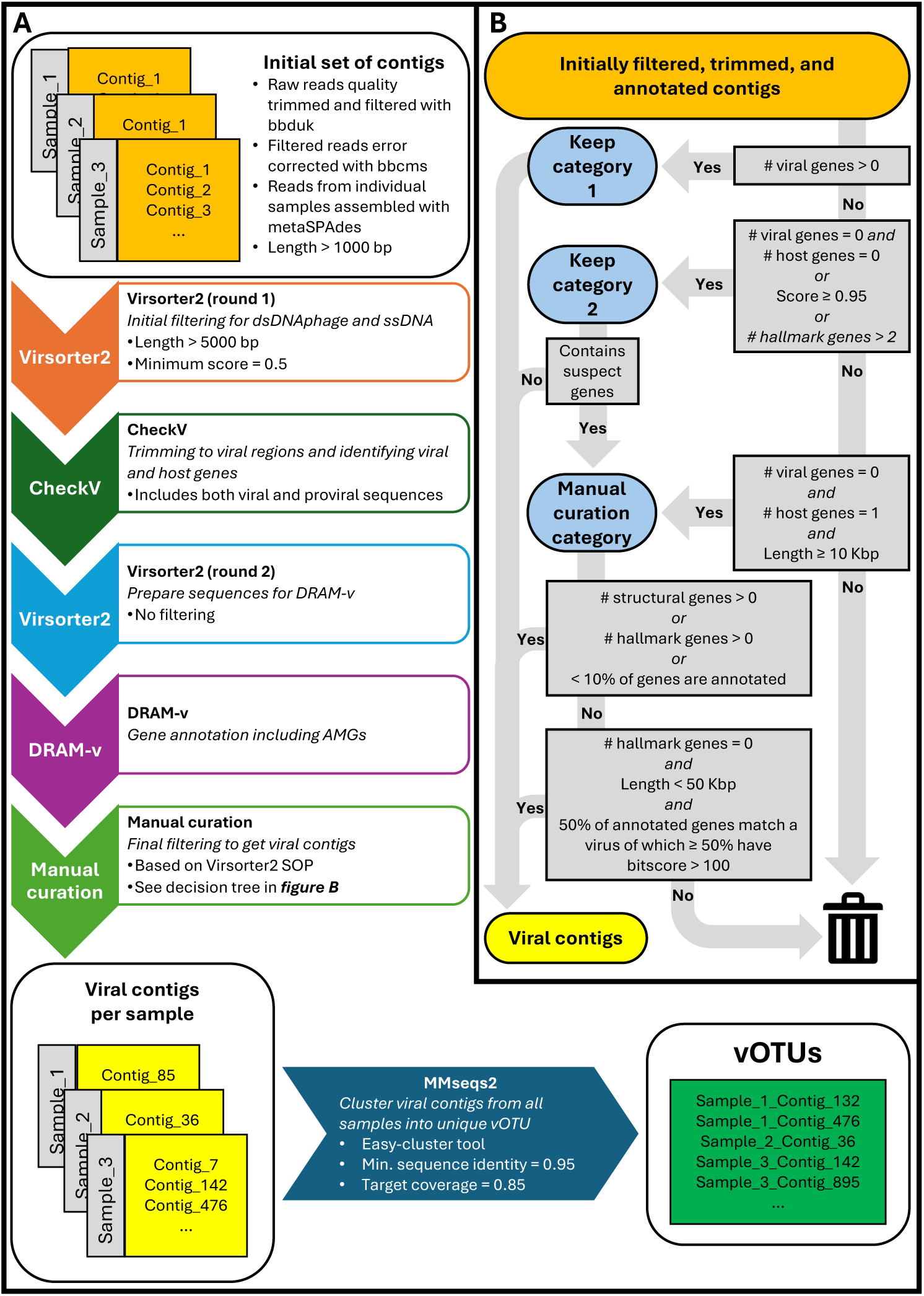
Overview of bioinformatic sequence processing used in this study. A) Flowchart describing the overall processing of assembled contigs into vOTU. B) Decision tree of how initially filtered, trimmed, and annotated contigs are filtered into viral contigs.

Reads across all samples were mapped to vOTUs using BBmap (Bushnell, 2019). vOTU were clustered with the INPHARED reference database (downloaded November 9^th^, 2023) (Cook et al., 2021) using vContact2 (Bin Jang et al., 2019) and taxonomy assigned using graphanalyzer (Mattia et al., 2022). vOTU quality was determined using CheckV (Nayfach et al., 2021). In an attempt to match vOTU to potential hosts, CRISPR spacers, identified from all contigs with minCED version 0.3.2 (https://github.com/ctSkennerton/minced), were aligned to vOTU using BLASTn (Altschul et al., 1990).

### 2.4 Statistical analysis

All statistical analyses were performed in R version 4.2.2 (R Core Team, 2018) with code available via GitHub (https://github.com/ShadeLab/Centralia_phages_Barnett). Since our experimental design utilized repeated sampling of sites over seven years, we used linear mixed effects (LME) models from package nlme (Pinheiro et al., 2020) to compare count, diversity, and abundance measures across time and soil temperature. For all LME models, the site was the random effect. Differences in measures between fire classification (fire-affected versus reference soils) were determined using the nonparametric Wilcoxon rank sum test. We used the Benjamini-Hochburg procedure for all post-hoc tests to adjust p-values for multiple comparisons.

vOTU were determined to be present in a sample if the average fold coverage was ≥ 1 and the percent of the vOTU sequence covered was ≥ 75%. vOTU abundance was measured as reads per thousand base pairs per million mapped reads (RPKM). RPKM for each vOTU was calculated for each sample as the number of forward and reverse reads mapped to the vOTU in that sample divided by the length of the vOTU in Kbp and then further divided by the total number of reads mapped to all vOTU in that sample divided by 1,000,000. Between sample diversity (beta diversity) was calculated with the Bray-Curtis dissimilarity index through package vegan (Oksanen et al., 2018). Variation in beta diversity was explained by fire classification and year, and their interaction was determined using PERMANOVA through package vegan with samples blocked by site ID to account for repeated sampling. To see if phage community beta-diversity differed between spatial and temporal scales, we compared the Bray-Curtis dissimilarities between sites within each year to dissimilarities within each site over time using the Wilcoxon rank sum test. We further performed beta diversity partitioning on these diversity measures to distinguish the influence of balanced variation in species abundances from abundance gradients in vOTU community dissimilarity (Baselga, 2013). Beta diversity portioning was performed using package betapart (Baselga and Orme, 2012) and compared across spatial and temporal scales using the Wilcoxon rank sum test. Community directionality over time was calculated from Bray-Curtis dissimilarity measures using the package ecotraj (De Cáceres et al., 2019; Sturbois et al., 2021). We tested whether the dissimilarity in community structure between sites within a year was different from dissimilarity over time within a site using the Wilcoxon rank sum test.

Finally, we wanted to compare the beta diversity of the viral community (vOTUs) to that of the bacterial communities (OTU). Bacterial OTU were processed as previously described (Barnett and Shade, 2024a). As the previous study had more sites than included here, we removed samples not included in metagenome sequencing and rarefied the resulting OTU table to the lowest sequencing depth (161,171 reads). Bray-Curtis dissimilarity in bacterial communities was calculated as before. To compare viral and bacterial community beta diversity structures, we performed a Mantel test using the two Bray-Curtis dissimilarity matrices and a Procrustes analysis using the principal coordinate analysis ordinations (PCoA) of the two Bray-Curtis dissimilarity matrices. After scaling and centering, we also performed a partial Mantel test, controlling for the Euclidian distance in both pH and soil temperature between samples to remove any potential correlation due to these notable edaphic factors (Barnett and Shade, 2024a). Mantel tests, PCoA ordination, and Procrustes analysis were all performed using the vegan package.

## 3 Results

### 3.1 Bacterial investment in CRISPR arrays increased with disturbance intensity

The metagenome investment in CRISPR arrays is a signature of potential phage-host interactions. Investment in CRISPR arrays was measured as the number of CRISPR arrays identified relative to the median number of copies of ribosomal proteins found in a sample. Overall, we found that the investment in CRISPR arrays was higher in the fire-affected sites than in the reference sites (Fig 2A; Wilcoxon test: W = 49, p-value < 0.001). Regardless of fire classification, CRISPR array investment increased with soil temperature (Fig 2B; LME: p-value < 0.001). Investment in CRISPR decreased over the seven years in the fire-affected sites but not reference sites (Fig fire-affected LME: p-value < 0.001, reference LME: p-value = 0.165).

**Figure 2:**
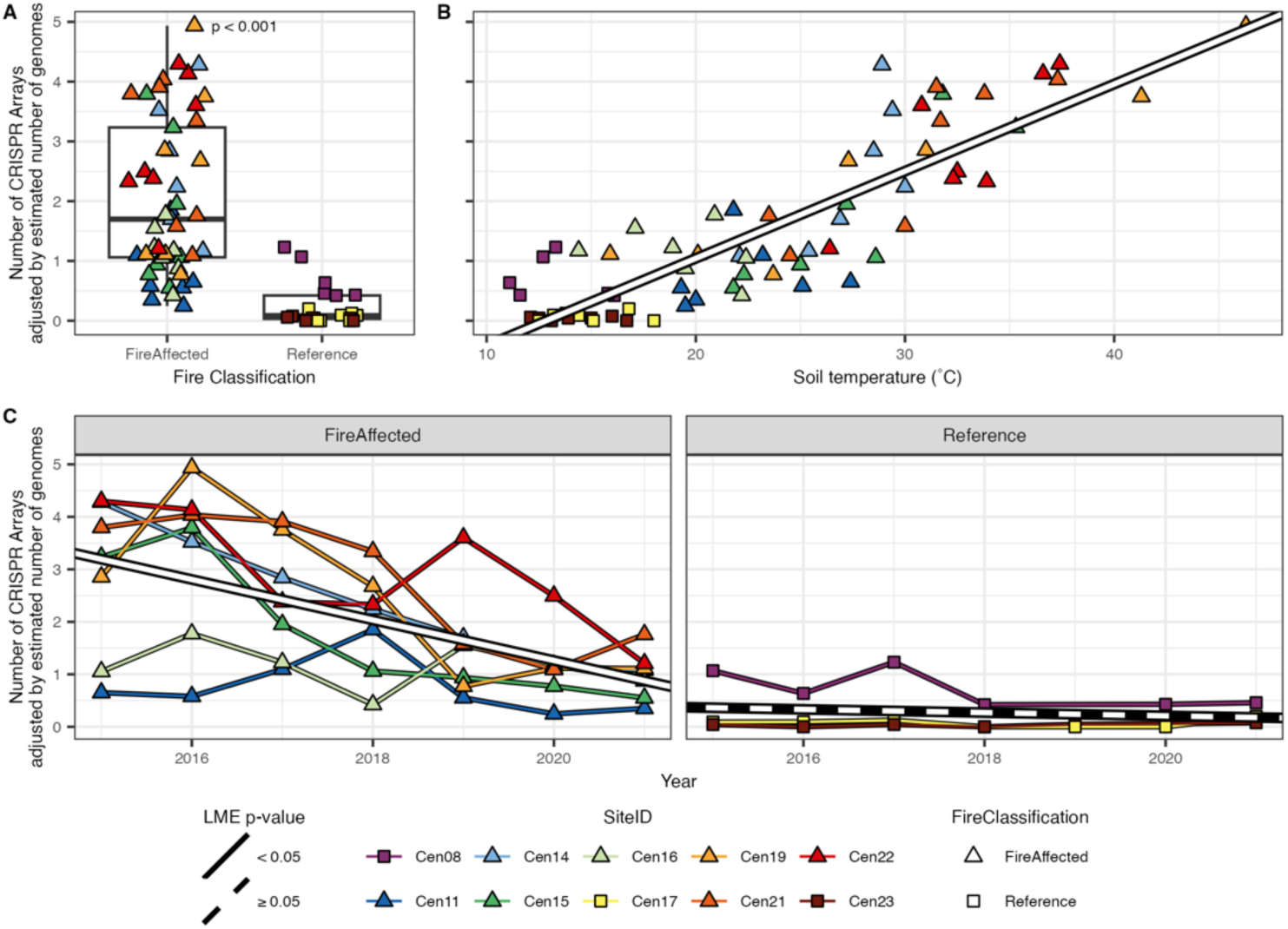
Community investment in CRISPR arrays is higher in fire-affected soils than in reference and decreases as these fire-affected soils cool. Community investment is calculated as the number of CRISPR arrays found divided by the median count of ribosomal protein within each sample. A) Compared across fire classification with the p-values of the Wilcoxon rank sum test indicated. B) Compared across soil temperature (°C) regardless of fire classification with the line indicating LME regression (p-value < 0.001). C) Compared across sampling years within fire classification with the line indicating LME regression (fire-affected p-value < 0.001, reference p-value = 0.165).

### 3.2 vOTU diversity varied with soil temperature and time

We found a total of 6929 vOTU across all Centralia soils. To assess within-sample vOTU diversity, we used two main measures: the percentage of sequenced reads mapped to vOTU (Fig. S2), and richness adjusted by total number of reads mapped to vOTU (Fig. S3). More reads mapped to vOTU in the fire-affected soils than reference soils (Wilcoxon test: W = 42, p-value < 0.001), and the percentage of reads mapping to vOTU decreased over time in fire-affected soils (LME: p-value < 0.001) but not reference soils (LME: p-value = 0.818). Correspondingly, the percentage of reads mapping to vOTU decreased with decreasing temperature (LME: p-value < 0.001). However, relatively fewer vOTU were found in the fire-affected soils than in reference soils (Wilcoxon test: W = 898, p-value < 0.001), and vOTU richness increased over time in fire-affected soils (LME: p-value < 0.001) but not in reference soils (LME: p-value = 0.358). Correspondingly, vOTU richness increased with decreasing temperature (LME: p-value < 0.001). Next, we used Pielou’s evenness, using RPKM as a proxy of abundance, to examine the patterns in relative abundances of vOTU within the soils (Fig. 3A-C). vOTU abundances were less even in the fire-affected soils than in the reference soils (Wilcoxon test: W = 874, p-value < 0.001), and vOTU evenness increased over time in fire-affected soils (LME: p-value < 0.001) but not reference soils (LME: p-value = 0.875). Correspondingly, vOTU evenness increased with decreasing temperature regardless of fire classification (LME: p-value < 0.001). While new unique vOTUs were detected in each consecutive year, particularly in fire-affected sites (Fig. 3D), within-site vOTU accumulation approached saturation within each site over time, suggesting low cross-year variability within a site. However, when considering each additional site sampled within any single year, there was continued and sharp increase in vOTU accumulation, suggesting high site-to-site variability within a year (Fig. 3E).

**Figure 3:**
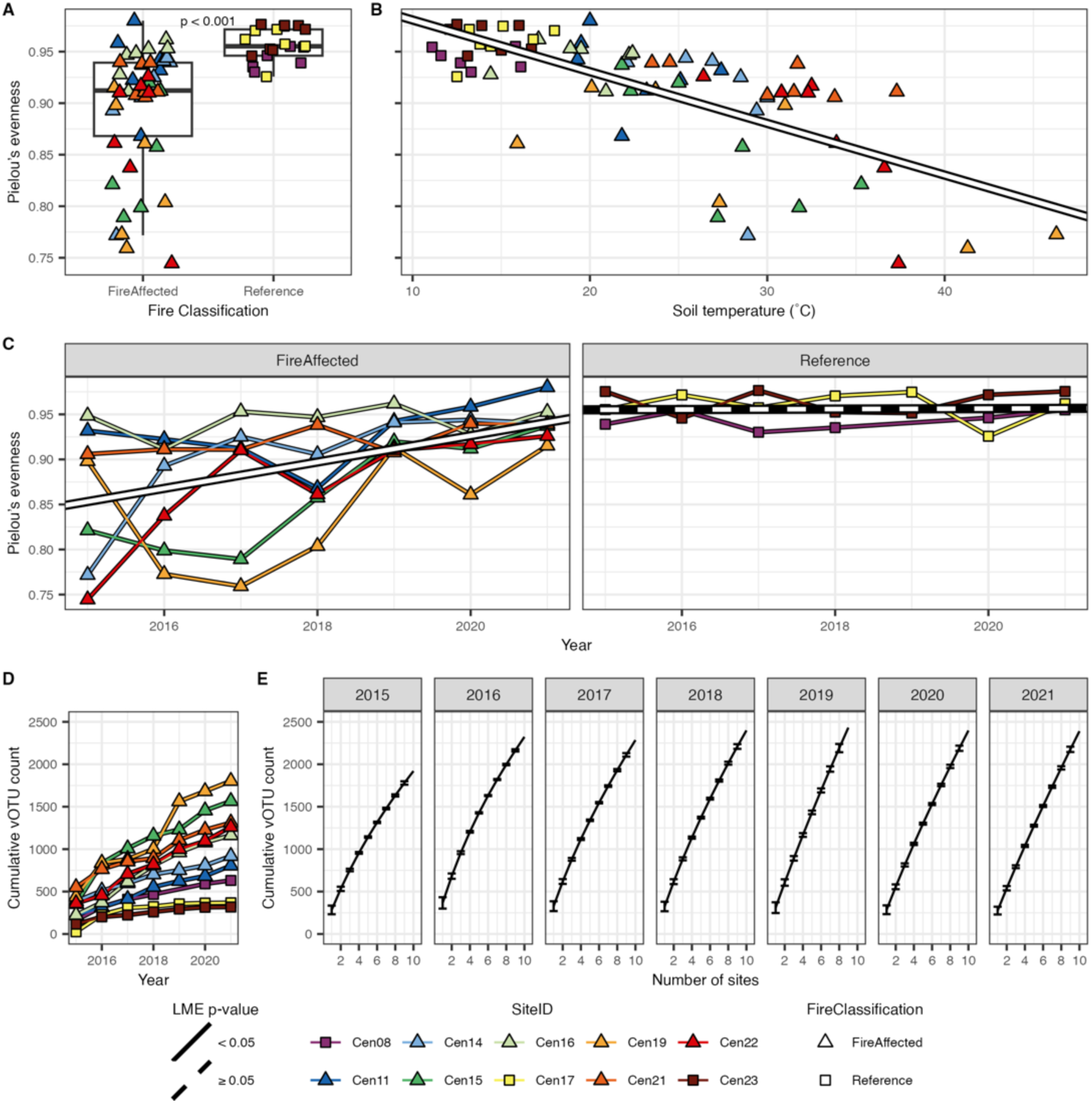
vOTU community evenness is higher in fire-affected soils than reference and decreases as these fire-affected soils cool. vOTU community evenness is measured as Pielou’s evenness. A) Compared across fire classification with the p-values of the Wilcoxon rank sum test indicated. B) Compared across soil temperature (°C) regardless of fire classification with the line indicating LME regression (p-value < 0.001). C) Compared across sampling years within fire classification with the line indicating LME regression (fire affected p-value < 0.001, reference p-value = 0.875). D) Cumulative vOTUs were uncovered at each site during each successive sampling year. C) Cumulative vOTUs were uncovered with additional site sampling within each sampling year, with the line indicating mean unique vOTU count and error bars indicating standard error across all sets of sites.

To examine across-sample vOTU diversity, we used Bray-Curtis dissimilarity with RPKM to measure abundance. Variation in vOTU community structure across samples (Fig. 4A) was explained by fire classification (R^2^ = 0.086; p-value = 0.002), time (R^2^ = 0.064; p-value = 0.001), and their interaction (R^2^ = 0.060; p-value = 0.005). Unlike the bacterial OTU diversity (Barnett and Shade, 2024a), there was no variation in beta dispersion across fire classification for vOTU (p-value = 0.516). We found that time only explains variation across fire-affected samples (post-hoc PERMANOVA R^2^ = 0.102, adjusted p-value = 0.006) but not across reference samples (adjusted p-value = 0.404). In agreement with the vOTU accumulation curves, vOTU dissimilarity varied more across sites within a year (*i.e.,* spatial scale) than within a site across years (*i.e.,* temporal scale) (Fig. 4B; Wilcoxon-test; W = 57064, p-value < 0.001). Upon partitioning these dissimilarities (Baselga, 2013), we observed greater abundance gradients (overlap and gradual changes in taxa along a gradient) within a site across years than across sites (Fig. 4C; Wilcoxon-test; W = 7557, p-value < 0.001) but greater balanced variation in species abundances (replacement of taxa along a gradient) across sites than within a site (Fig. 4D; Wilcoxon-test; W = 57210, p-value < 0.001).

**Figure 4:**
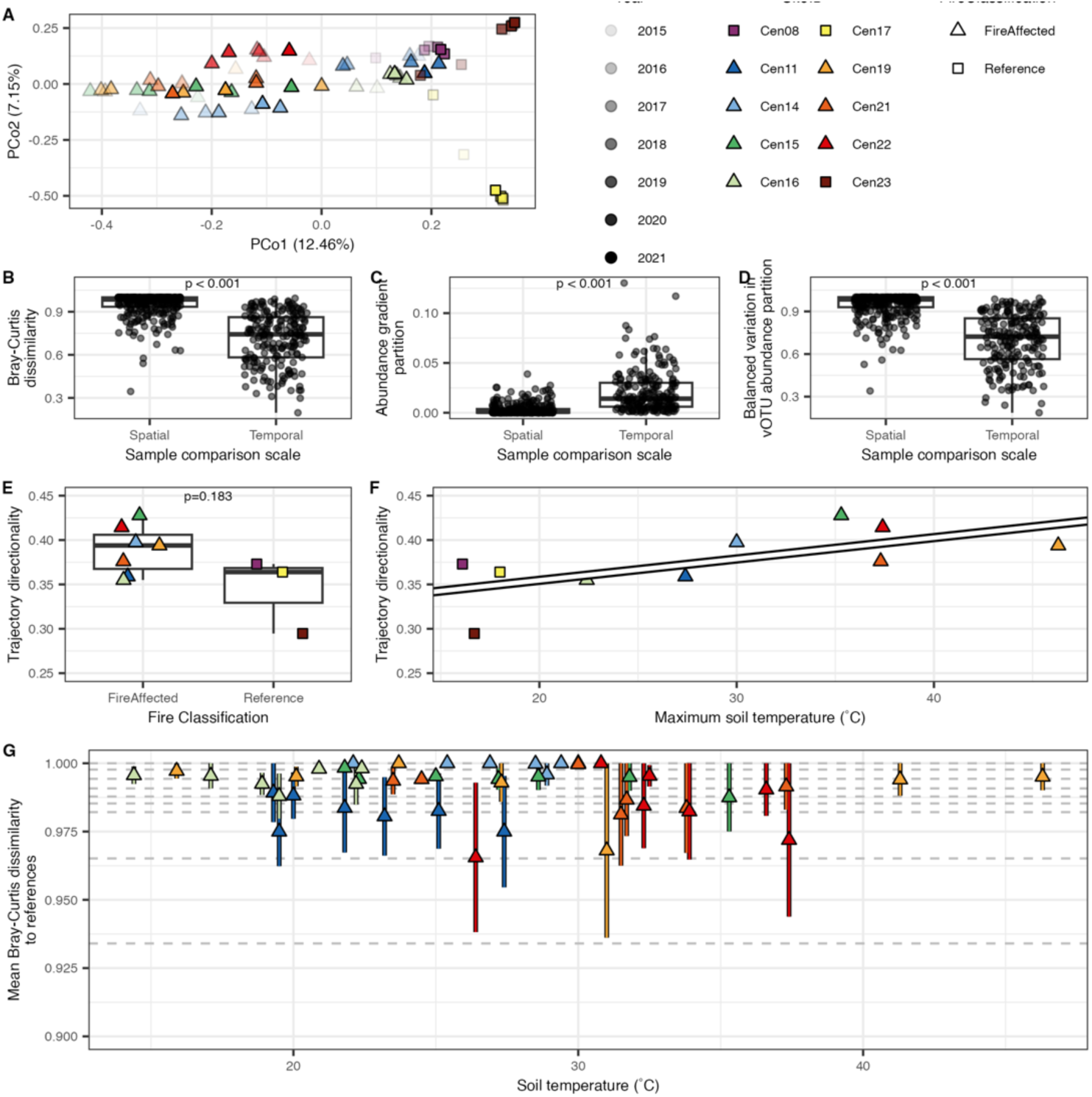
Variation in vOTU diversity, as measured by Bray-Curtis dissimilarity, is explained by fire classification and sampling year. A) Principal coordinate analysis ordination with site ID, fire class, and sampling year indicated. Axis labels give the percent of the variation in vOTU diversity explained by each axis. B) Bray-Curtis dissimilarity across samples differing spatially (*i.e.,* different sites but same sampling year) or temporally (*i.e.,* same site but different sampling years). C) Partition of Bray-Curtis dissimilarity is explained by an abundance gradient across samples differing spatially or temporally, similar to community nestedness. D) Partition of Bray-Curtis dissimilarity explained by balanced variation in vOTU abundance across samples differing spatially or temporally, similar to community turnover. E) Community directionality for each site over time across fire classification. F) Community directionality for all sites regardless of fire classification against maximum soil temperature. The solid line represents the LME regression (p-value = 0.034). G) Mean community dissimilarity between each fire-affected site and the three reference sites across fire-affected soil temperature in the corresponding sampling year. Error bars indicate standard error (n=3). Dashed lines indicate the community dissimilarity between the reference sites within sampling years. There was no statistically significant relationship across dissimilarity between fire-affected and reference soil and soil temperature (LME: p-value = 0.721). P-values included in the figures indicate the Wilcoxon rank sum tests.

Trajectory analysis, which examines shifts in community composition over time, found no statistically significant variation in viral community directionality between fire-affected and reference communities on the whole (Fig. 4E; Wilcoxon-test; W = 17, p-value = 0.183), but directionality was positively associated with maximum soil temperature at each site (*i.e.,* disturbance intensity; Fig. 4F; linear regression; R^2^ = 0.379, p-value = 0.034). However, there was no change in Bray-Curtis dissimilarity between fire-affected and reference sites as the soils cooled (Fig. 4G; linear mixed effects model; p-value = 0.721), likely due to exceedingly high dissimilarity in viral community composition (Bray-Curtis dissimilarity > 0.9) across all sites at all time points.

### 3.3 Dissimilarity in viral community structure is correlated to bacterial community structure

vOTU community structure (Bray-Curtis dissimilarity) correlated strongly with bacterial structure (Mantel test: r = 0.5113, p-value < 0.001). There was also concordance between vOTU and bacterial OTU community structures across samples (Procrustes: m12 squared = 0.1502, p-value = 0.001). After accounting for correlation explained by soil temperature and pH, vOTU and bacterial community structures were more weakly correlated (partial Mantel test: r = 0.3863, p-value < 0.001). Notably, bacterial community structure was highly correlated to differences in soil temperature and pH (Mantel test: r = 0.6717, p-value < 0.001), as previously shown (Barnett and Shade, 2024a),.

### 3.4 vOTU taxonomy was dominated by unknown viruses

Over 90% of the vOTUs were unclassified, with another 8.6% classified in the Caudoviricetes class but unclassified at the family level. Of the other classified taxa, in the *Caudoviricetes* class there were nine *Autographiviridae*, five *Steigviridae*, three *Mesyanzhinovviridae* and *Winoviridae*, two *Herelleviridae* and *Peduoviridae*, and one each from *Anaerodiviridae, Kyanoviridae, Podoviridae, Pungoviridae,* and *Zobellviridae*. We also found 27 *Tectiviridae* from class *Tectiliviricetes*, two *Inoviridae* from class *Faserviricetes*, and one *Microviridae* from class *Malgrandaviricetes*.

### 3.5 Few vOTUs had matching CRISPRs

There were 184 vOTU (2.66% of vOTU) that matched one or more CRISPR spacers across samples. These CRISPR spacers (2672 spacers) represented 2.46% of all identified CRISPR spacers across samples but 10.48% of all CRISPR arrays (674 arrays). Over half of these CRISPRs (1726 spacers and 425 arrays) were detected in the same samples as their mapped vOTUs, indicating the co-occurrence of many of these host-phage pairings (Fig. S4). Notably, at least one vOTU-CRISPR spacer co-occurrence was observed in all seven fire-affected sites, but there were no such co-occurrences detected in any reference site. While over half of CRISPR arrays that matched at least one vOTU were present on contigs unclassified at the phylum level (366), 120 were classified in the phylum *Pseudomonadota*, 86 were classified as *Actinomycetota*, 44 were classified as *Planctomycetota*, and less than ten each were classified in *Cyanobacteriota, Verrucomicrobiota, Acidobacteriota, Bacillota, Bacteroidota, Myxococcota, Deinococcota, Euryarchaeota, Thermomicrobiota, Chloroflexota,* and *Thermotogota*.

### 3.6 Few vOTU had annotated auxiliary metabolic genes

Out of all vOTU, we found 790 putative auxiliary metabolic genes (AMGs) in 533 vOTU (7.69% of vOTU). Considering all AMGs (Fig. S5), the cumulative abundance of vOTU-containing an annotated AMG did not vary across fire-affected and reference sites (Wilcoxon test: W = 491, p-value = 0.995), soil temperature (LME: p-value = 0.581), or time (LME: fire affected p-value = 0.713, reference p-value = 0.951). When examining only AMGs annotated as carbohydrate-active enzyme genes (CAZymes), the cumulative abundance of vOTU-containing CAZymes was higher in fire-affected sites than the reference sites (Fig. 5A; Wilcoxon test: W = 192, p-value < 0.001) and decreased with decreasing temperature (Fig. 5B; LME: p-value = 0.040). However, while the cumulative abundance of vOTU-containing peptidase annotated AMGs was also high in the fire-affected sites (Fig. 5C; Wilcoxon test: W = 192, p-value < 0.001), there was no relationship with temperature (Fig. 5D; LME: p-value = 0.791).

**Figure 5:**
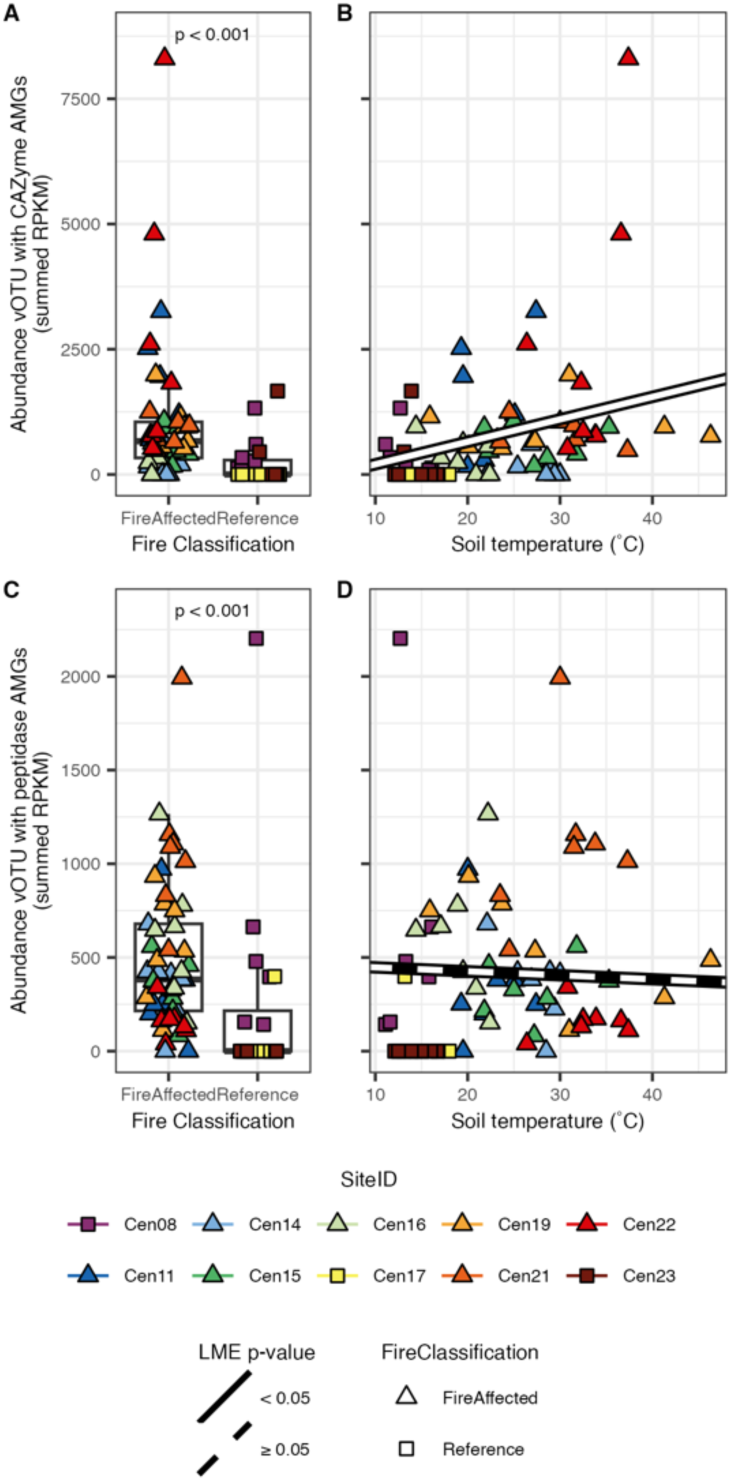
Abundance of vOTU encoding annotated CAZyme and peptidase AMGs are higher in fire-affected soils than reference soils, and there is a positive relationship between CAZyme encoding vOTU abundance and soil temperature. Abundances are measured as summed RPKM. CAZyme encoding vOTU abundance were compared A) between fire classifications and B) across soil temperature. Peptidase encoding vOTU abundance were compared C) between fire classifications and D) across soil temperature classifications. The Wilcoxon rank sum test between fire classifications is indicated above the points for box plots. For scatter plots, the LME regression is plotted as a solid (p-value < 0.05) or dashed (p-value ≥ 0.05) line.

## 4 Discussion

We examined seven years of viral diversity dynamics in soils associated with the Centralia, Pennsylvania, coal mine fire. We found high site-to-site variation in viral community structure, as previously well reported (Noah et al., 2007; Srinivasiah et al., 2015; Emerson et al., 2018; Trubl et al., 2018; Chevallereau et al., 2022; Durham et al., 2022; Santos-Medellín et al., 2022; Barnett and Buckley, 2023; Graham et al., 2023), but surprisingly consistent viral communities within a site across the seven years, with the reference soils having a relatively more stable community that the fire-affected. Furthermore, we found that, while the bacterial and viral communities were correlated to each other and to the major environmental drivers of temperature and pH, the viral communities did not exhibit the same degree of resilience and recovery to the fire as the bacterial communities.

It has been suggested that the high spatial variability in soil viral communities, even at small scales (*i.e.,* meters), is due to strong dispersal limitation (Santos-Medellín et al., 2022). Our finding of greater abundance variation and lower balanced variation contributions to vOTU dissimilarity within sites over time relative to across sites within a timepoint supports this hypothesis. There were fewer unique vOTU discovered with successive annual sampling within a site as compared to successive site sampling within a year. In other words, the composition of the vOTUs within a site did not change as much over time, despite dynamic environmental conditions within and across years. This lack of viral community turnover may be due to limited immigration of new phages into a site even over the years, with viral community changes primarily attributed to variations in the abundances of persistent members. As an expected limitation in using metagenome methods to understand viral communities (e.g., due to imperfect assembly and annotation methods particularly for viruses), some turnover observed in Centralia may be due to the recovery of previously unassembled viral sequences. Regardless, these results were robust across the sites included in this study, which represented a broad range of environmental contexts.

While we previously observed strong resilience in the bacterial communities as the temperatures cooled with fire advancement in Centralia (Barnett and Shade, 2024a), there was less evidence for viral community resilience. Over time, and even as soils cooled, highly dissimilar phage communities remained across fire-affected and reference communities, likely due to the inherently high spatial variability in viral structure across sites. While viral community directionality was highest in the soils undergoing the hottest recorded temperature over the seven sampling years, there was no overall difference in directionality between the viral communities across fire-affected and reference soils. These results suggest that the phage community composition remains distinct across disturbed soils for a longer duration than that of the bacterial communities. Phage diversity has even been shown to be uncoupled from microbial diversity in some soil systems (Ma et al., 2024). Thus, changes in bacterial communities during recovery may not lead to comparable changes in their phage communities. While both viral and bacterial community dissimilarity across samples were correlated, this correlation was reduced when controlling for correlation with soil temperature and pH. This uncoupling of phage and host communities may be partly due to the previously suggested limited dispersal of viruses across local spatial scales.

Our results suggest that there could have been a change in the interactions between phage and bacterial hosts across the disturbance gradient and during soil recovery. CRISPR investment and phage community evenness shifted as soils cooled, with both measures becoming more like those of undisturbed reference sites. Uneven phage communities suggest that there were a few highly abundant phages, typical of predator-prey dynamics such as “kill-the-winner” (Rodriguez-Brito et al., 2010). It is possible that Centralia fire-affected soils may be under more intense predator-prey relationships than undisturbed soils, possibly due to an observed decrease in bacterial community richness in the hotter sites (e.g., fewer host taxa that have relatively more phage infections). Increased phage infection and bacterial biomass turnover has previously been observed in experimental soils incubated at 23°C relative to 18°C incubations leading to increased soil carbon mineralization (Wang et al., 2022). High temperatures may also induce temperate phages to become lytic, increasing host population turnover (Jansson and Wu, 2022).

We observed more CRISPR arrays in the fire-affected soils than in the reference soils. CRISPR mechanisms are used by bacteria and archaea to combat phage infection through targeted and adaptive destruction of viral DNA. Overall, CRISPR-Cas abundance is correlated with viral abundance across many ecosystems, linking CRISPR system investment with environmental viral diversity (Meaden et al., 2022). Previous work that examined metagenomes from eight years of experimental warming of soils to about 4°C above ambient demonstrated an increased prevalence of the CRISPR-Cas marker gene, *cas1*, among populations of diverse phyla. In that study, increased prevalence was defined as the abundance of *cas1* genes normalized to the abundance of the phylum they were classified to. Such increased prevalence suggested selection for phage defense in warmed soil (Wu et al., 2020). Our study used a comparable measure of CRISPR investment by normalizing CRISPR array counts with estimated genome counts determined by median ribosomal protein gene counts, regardless of taxonomy. Our findings thus agree with this previous work despite differences in warming intensity (generally over 10°C above ambient temperature in Centralia) and determination of genomic investment/prevalence.

While we did not find many auxiliary metabolic genes (AMG) in the vOTUs detected and annotated, the AMGs we found suggest that these phages may influence bacterial metabolisms associated with ecosystem functioning (Breitbart et al., 2007). AMGs expressed in infected bacterial cells can reprogram the cellular metabolism and cellular function in ecosystem processes such as carbon and nitrogen cycling (Breitbart, 2011; Hurwitz et al., 2013). In soils, in particular, AMGs are believed to be important in carbon processing by providing hosts with new or alternative genes for polysaccharide binding, breakdown of complex carbohydrates, and central carbon metabolism (Trubl et al., 2018). Across the Centralia soils, while there was not any variation in the abundance of vOTUs containing one or more annotated AMGs across fire classifications and temperature, there was a higher abundance of vOTU with CAZyme and peptidase AMGs in fire-affected soils relative to reference soils. There was also a positive linear relationship between the abundance of vOTU-encoding CAZyme AMGs and soil temperature. Therefore, given their relatively higher abundance in warmer Centralia soils, vOTU may indirectly influence carbohydrate and protein metabolisms within hotter, more disturbed soils than undisturbed soils. While a limitation of this work is, like that of other studies using untargeted metagenome approaches to assay viral communities, the under-sampling of the viral community and incomplete annotations of AMG, our results demonstrate that disturbances may influence the activity of soil phages in ecosystem functioning beyond the direct response to the changes in their host populations.

## 5 Conclusion

Anthropogenic press disturbances result in large changes in soil microbial communities, including viruses. While we observed that soil heating from the coal seam fire underneath Centralia, Pennsylvania, has driven large-scale changes in bacterial and phage communities, there was evidence for relatively less recovery in the phage communities than we previously observed for the bacteria. Notably, there was less variation in soil viral communities over years within a site than across sites within a year, suggesting some consistency in viral community structure over time. Overall, this work provides insights into how viral communities respond to warming disturbance and how they change interannually, with both dynamics compared relative to those of their host communities. This work is relevant for understanding and predicting microbial responses to the changing climate.

## Acknowledgments

We thank Tammy Tobin for Centralia research consultation and local sampling support and Keara L Grady for coordinating fieldwork logistics and organization of multi-year soil procurement and processing. We additionally thank SH Lee, JW Sorensen, TK Dunivin, JL Chodkowski, N Stopnisek, and M Mechan Llontop for collecting samples. This work was supported by the U.S. National Science Foundation CAREER award #1749544 to AS. AS acknowledges the French Centre National de la Recherche Scientifique (CNRS) support. We declare no conflicts of interest.

