## Supplemental for "Soil viral community dynamics over seven years of heat disturbance: spatial variation exceeds temporal in annually sampled soils"

**Supplemental Tables and Figures**

| SiteID | Fire Classification | Temp. 2015 (˚C) | Temp. 2016 (˚C) | Temp. 2017 (˚C) | Temp. 2018 (˚C) | Temp. 2019 (˚C) | Temp. 2020 (˚C) | Temp. 2021 (˚C) | Latitude and Longitude |
|---|---|---|---|---|---|---|---|---|---|
| Cen08 | Reference | 12.7 | 11.1 | 13.3 | 16.1 | 16.3 | 11.6 | 15.8 | 40.8014N 76.3461W |
| Cen11 | FireAffected | 27.4 | 25.1 | 23.2 | 21.8 | 19.3 | 19.5 | 20 | 40.8013N 76.3431W |
| Cen14 | FireAffected | 28.9 | 29.4 | 28.5 | 30 | 26.9 | 25.4 | 22.1 | 40.8007N 76.3412W |
| Cen15 | FireAffected | 35.3 | 31.8 | 27.2 | 28.6 | 25 | 22.3 | 21.8 | 40.8008N 76.3415W |
| Cen16 | FireAffected | 22.4 | 20.9 | 18.9 | 22.2 | 17.1 | 14.4 | 19.5 | 40.8008N 76.3414W |
| Cen17 | Reference | 14.4 | 13.2 | 13.8 | 18 | 15.1 | 12.5 | 16.8 | 40.8N 76.3403W |
| Cen19 | FireAffected | 31 | 46.3 | 41.3 | 27.3 | 23.7 | 15.9 | 20.1 | 40.8008N 76.3415W |
| Cen21 | FireAffected | 33.8 | 37.3 | 31.5 | 31.7 | 30 | 24.5 | 23.5 | 40.8007N 76.3414W |
| Cen22 | FireAffected | 37.4 | 36.6 | 32.3 | 33.9 | 30.8 | 32.5 | 26.4 | 40.8011N 76.3431W |
| Cen23 | Reference | 13.9 | 13.1 | 12.5 | 16.7 | 15 | 12.1 | 16 | 40.8013N 76.3466W |

**Table S1:** Geographical information on all 10 sites used in this study including fire classification, soil temperatures (Temp.) during each year, and site coordinates.

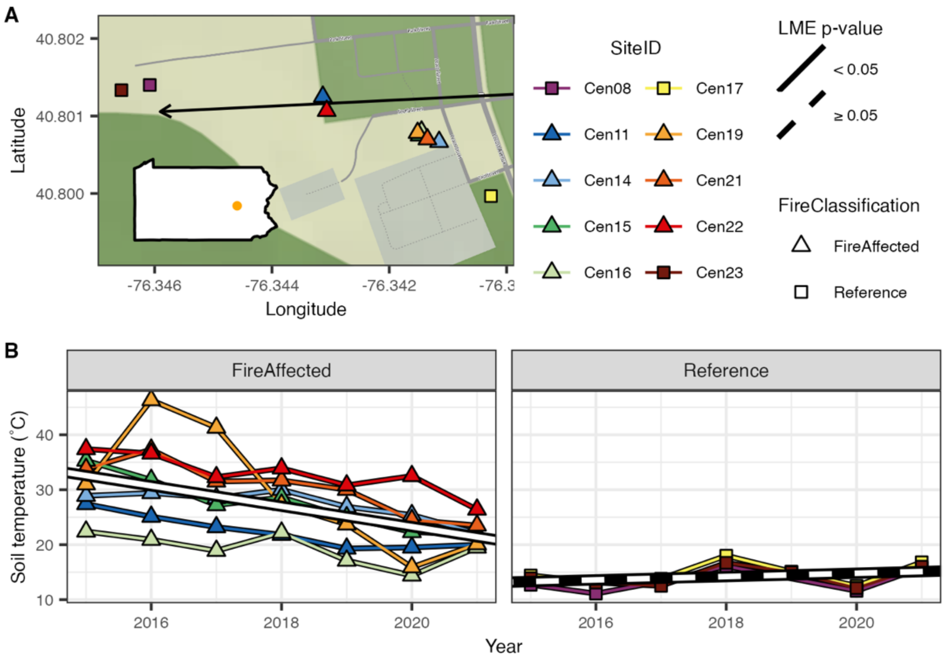


**Figure S1:** Seven fire affected and three reference sites in Centralia, Pennsylvania were sampled annually over seven years. A) map of Centralia with all 10 sites. Location of Centralia within Pennsylvania is included as intsert state map. B) Soil temperature within the fire affected sites decreased over the sampling years but there was no significant change in reference sites (LME: fire affected p-value < 0.001, reference p-value = 0.160).

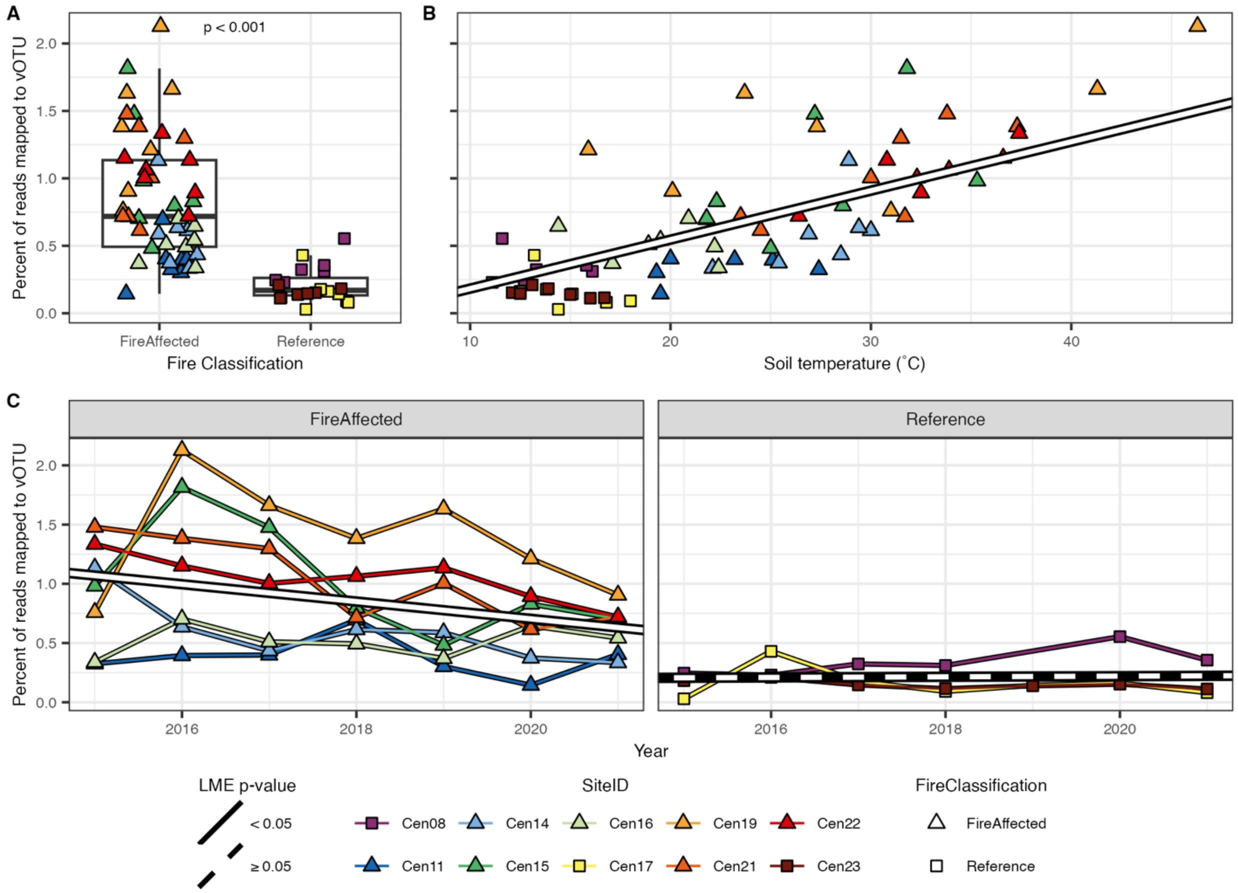


**Figure S2:** Percent of total reads mapping to vOTU is higher in fire affected soils than reference and decreases as these fire affected soils cool. A) Compared across fire classification with p-value of the Wilcoxon rank sum test indicated. B) Compared across soil temperature (˚C) regardless of fire classification with line indicating LME regression (p-value < 0.001). C) Compared across sampling years within fire classification with line indicating LME regression (fire affected p-value < 0.001, reference p-value = 0.818).

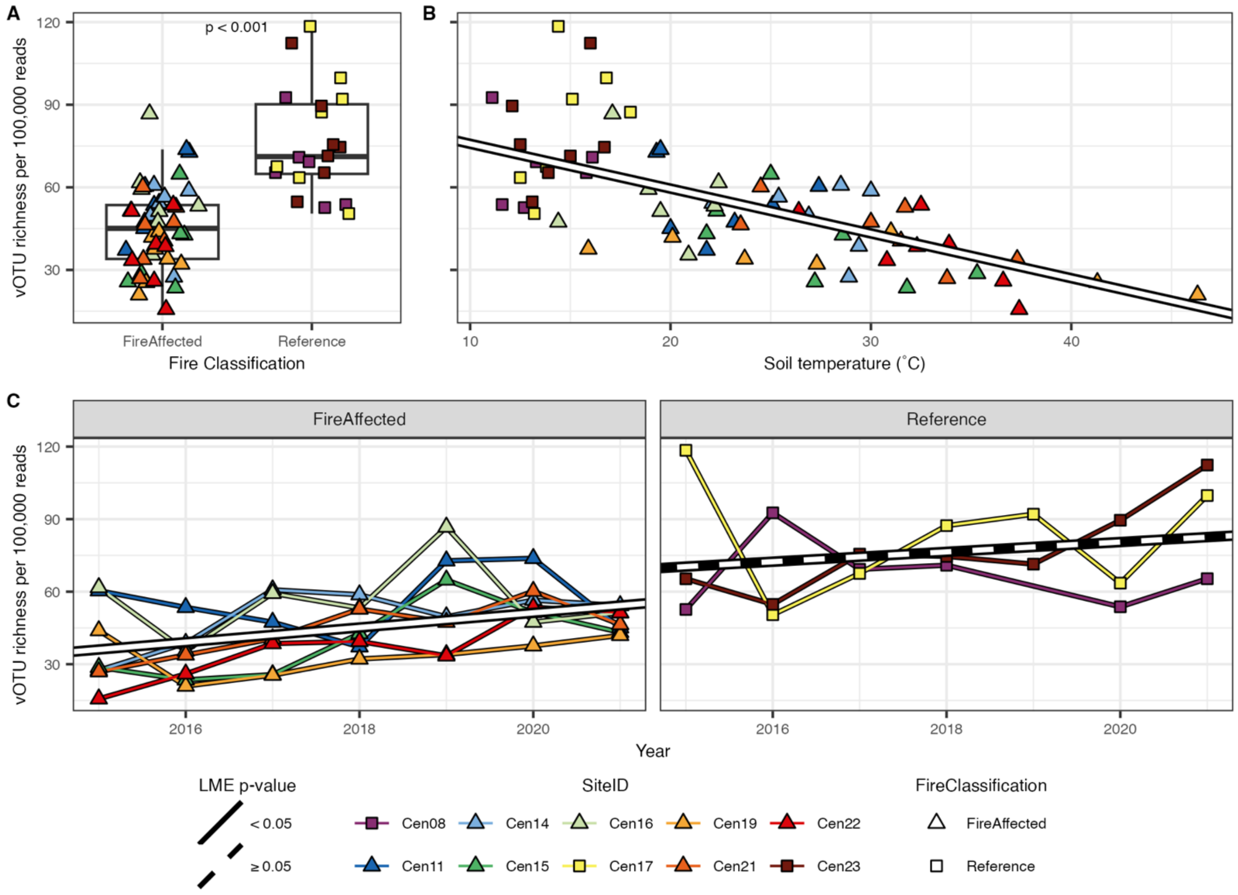


**Figure S3:** vOTU richness is higher in reference soils compared to fire affected and increases as the fire affected soils cool. vOTU richness is measured as number of vOTU observed per 100,000 reads mapped to all vOTU in the sample. A) Compared across fire classification with p-value of the Wilcoxon rank sum test indicated. B) Compared across soil temperature (˚C) regardless of fire classification with line indicating LME regression (p-value < 0.001). C) Compared across sampling years within fire classification with line indicating LME regression (fire affected p-value < 0.001, reference p-value = 0.358).

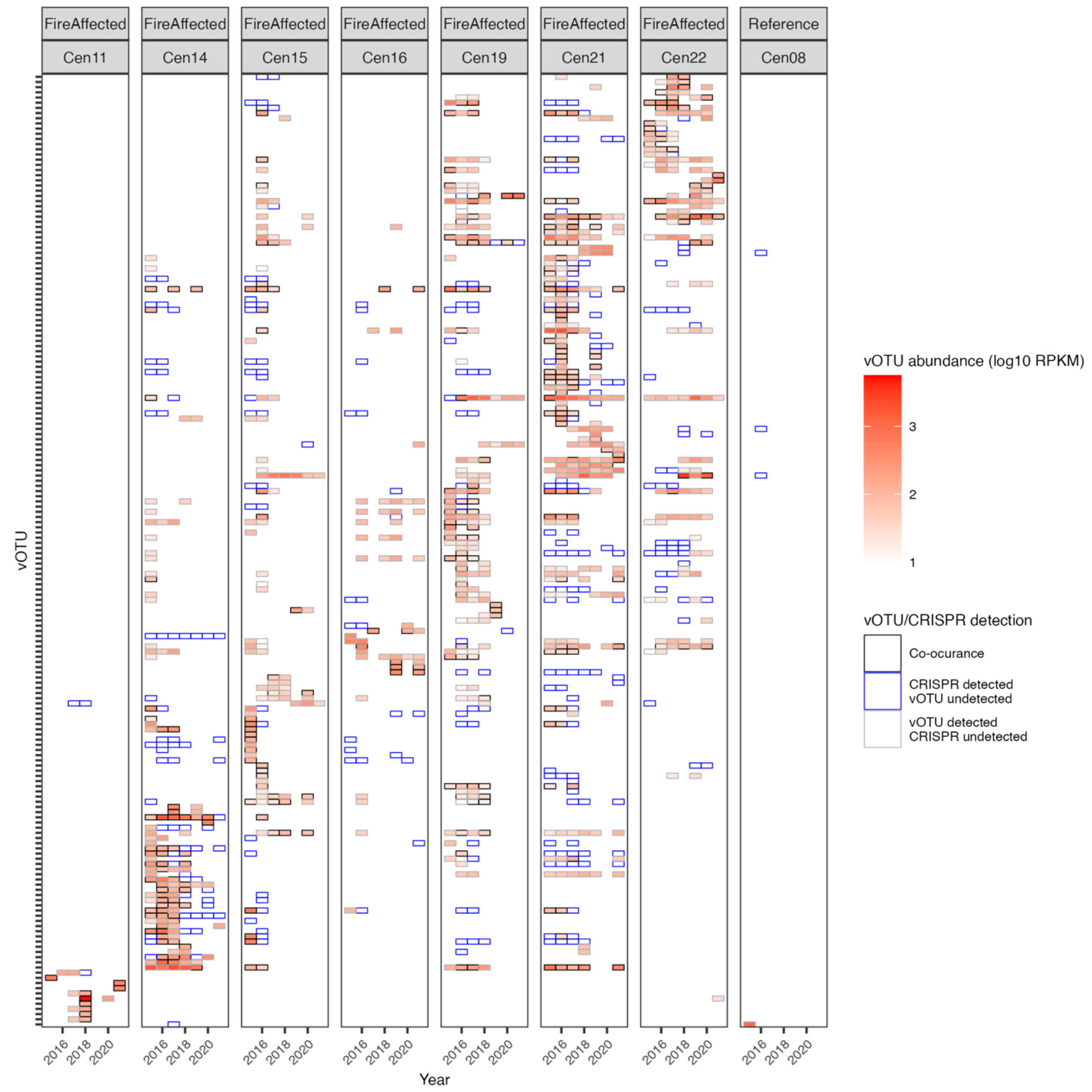


**Figure S4:** Heatmap of occupancy and abundance of vOTU matching CRISPR spacers across samples. Boxes represent vOTU detection (grey), CRISPR dectection (blue), or both detected (black). Red fill for vOTU detection and co-occurance indicates vOTU abundance in the sample ($\log_{10}$ RPKM). Only one reference site is included as no matching vOTU or CRISPRs were detected in the other two references (Cen17 and Cen23).

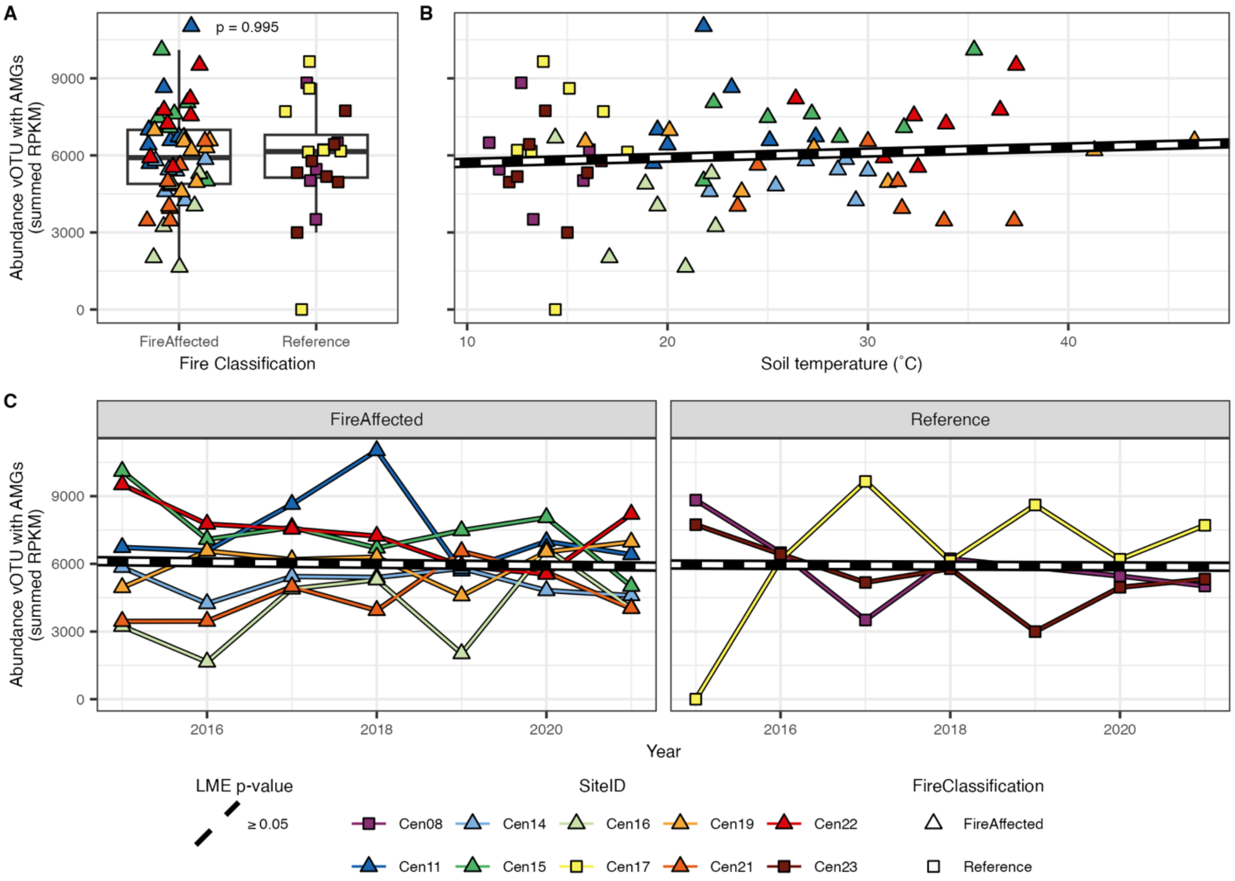


**Figure S5:** Abundance of vOTU encoding an annotated AMG did not significantly vary across fire classification, temperature or time. A) Compared across fire classification with p-value of the Wilcoxon rank sum test indicated. B) Compared across soil temperature (˚C) regardless of fire classification with line indicating LME regression (p-value = 0.581). C) Compared across sampling years within fire classification with line indicating LME regressions (fire affected p-value = 0.713, reference p-value = 0.951).
